# Intracellular helix-loop-helix domain modulates inactivation kinetics of mammalian TRPV5 and TRPV6 channels

**DOI:** 10.1101/2022.10.04.510840

**Authors:** Lisandra Flores-Aldama, Daniel Bustos, Deny Cabezas-Bratesco, Sebastián E. Brauchi

## Abstract

TRPV5 and TRPV6 are calcium-selective ion channels expressed at the apical membrane of epithelial cells. Important for systemic calcium (Ca^2+^) homeostasis, these channels are considered as gatekeepers of this cation transcellular transport. Intracellular Ca^2+^ exerts a negative control over the activity of these channels by promoting inactivation. TRPV5 and TRPV6 inactivation has been divided into fast and slow phases based on their kinetics. While slow inactivation is common to both channels, fast inactivation is characteristic of TRPV6. It has been proposed that the fast phase depends on Ca^2+^ binding and that the slow phase depends on the binding of the Ca^2+^/Calmodulin complex to the internal gate of the channels. Here, by means of structural analyses, site-directed mutagenesis, electrophysiology, and molecular dynamic simulations, we identified a specific set of amino acids and interactions that determine the inactivation kinetics of mammalian TRPV5 and TRPV6 channels. We propose that the association between the intracellular helix-loop-helix (HLH) and the TRP helix (TDh) domains favors the faster inactivation kinetics observed in mammalian TRPV6 channels.

## Introduction

Systemic calcium (Ca^2+^) homeostasis is vital for many physiological processes. Maintaining Ca^2+^ levels within a narrow range in extracellular fluids is regulated by the rates of absorption, storage, and excretion. These processes are mediated by passive and active transport mechanisms (1,2). Active transcellular transport is a regulated process involving three steps: apical entry of Ca^2+^ into epithelial cells, transport to the basolateral membrane, and extrusion to the bloodstream (1,2). This cellular mechanism is maintained by a specific set of transporters, pumps, and ion channels (1,2). TRPV5 and TRPV6 are calcium-selective ion channels belonging to the Transient Receptor Potential (TRP) family (3). Expressed at the apical membrane of calcium-transporting epithelia, these channels serve as entry channels in the transcellular pathway (1–3). Altered expression of TRPV5 and TRPV6 has been related to cancer (4–8), multiple pathologies of bone (9–13) and kidney (11,14–16), and impairments in reproductive physiology (17–19). In agreement with their physiological relevance, expression levels, trafficking, and activity of these channels are highly regulated (1).

An increase in the intracellular Ca^2+^ concentration induces fast and slow types of inactivation in calcium-selective TRP channels TRPV5 and TRPV6 (20). Slow inactivation is common to both, and it is determined by the binding of the Ca^2+^/Calmodulin complex to the channel’s intracellular region (21,22). Based on functional and structural data, it has been hypothesized that calmodulin (CaM) blocks the channel’s intracellular gate via a cation-π interaction between a lysine side chain from CaM (Lys116 in rat CaM) and tryptophan residues located at the lower gate of the channels (Trp583 in rabbit TRPV5). In addition to the slow inactivation component, a strong fast inactivation is observed only in mammalian TRPV6 channels (20,23). Our phylogenetic and functional studies suggested that calcium-dependent fast inactivation corresponds to an evolutive innovation in vertebrates that is absent in fish and nearly absent in reptiles (24).

Different groups have mapped residues associated with fast inactivation. It has been found that amino acids located at Helix-Loop-Helix (HLH) domain (residues E288, F292, and S298 in hTRPV5) (24), the intracellular linker between the transmembrane segments 2 and 3 (S2-S3 linker; residues L409, V411, T412, and F414 in hTRPV6) (23), and the lower gate (Q579 in hTRPV5, and H587 in hTRPV6) (25) form part of the mechanism of inactivation. In this context, our previous data suggest a strong evolutionary correlation between residues located at the HLH domain and the S2-S3 linker, supporting the robust inactivation displayed by mammalian TRPV6 channels (24).

In this study, we analyzed the sequence conservation of each element of the HLH/S2-S3 linker/TDh inactivation motif and identified a scaffold sequence at the HLH and S2-S3 linker harboring individual residues that are responsible for fast inactivation. By a close examination of the available three-dimensional structures of mammalian TRPV5 and TRPV6 channels in the apo and Ca^2+^/CaM bound states, we identified differences in the network of connectivity within this putative inactivation motif. Site-directed mutagenesis followed by patch-clamp electrophysiological recordings unveiled individual residues that are responsible for inactivation. Lastly, molecular dynamics simulations suggest that a condition where the HLH domain comes closer to the TDh is required to modulate fast inactivation of mammalian TRPV6.

## Results

### The evolutionary profiles of the inactivation motif

We first explored the elements of the HLH/S2-S3 linker/TDh motif by studying their primary sequences individually (Fig 1). The HLH and the S2-S3 linker display higher conservation in mammals and sauropsids and are evidently more variable in amphibians and fish (Fig. 1B). In contrast, the consensus sequence corresponding to the TDh is relatively conserved in all examined groups (Fig. 1B). This suggests independent evolutionary transitions for the different structural pieces forming the motif.

**Figure 1.**
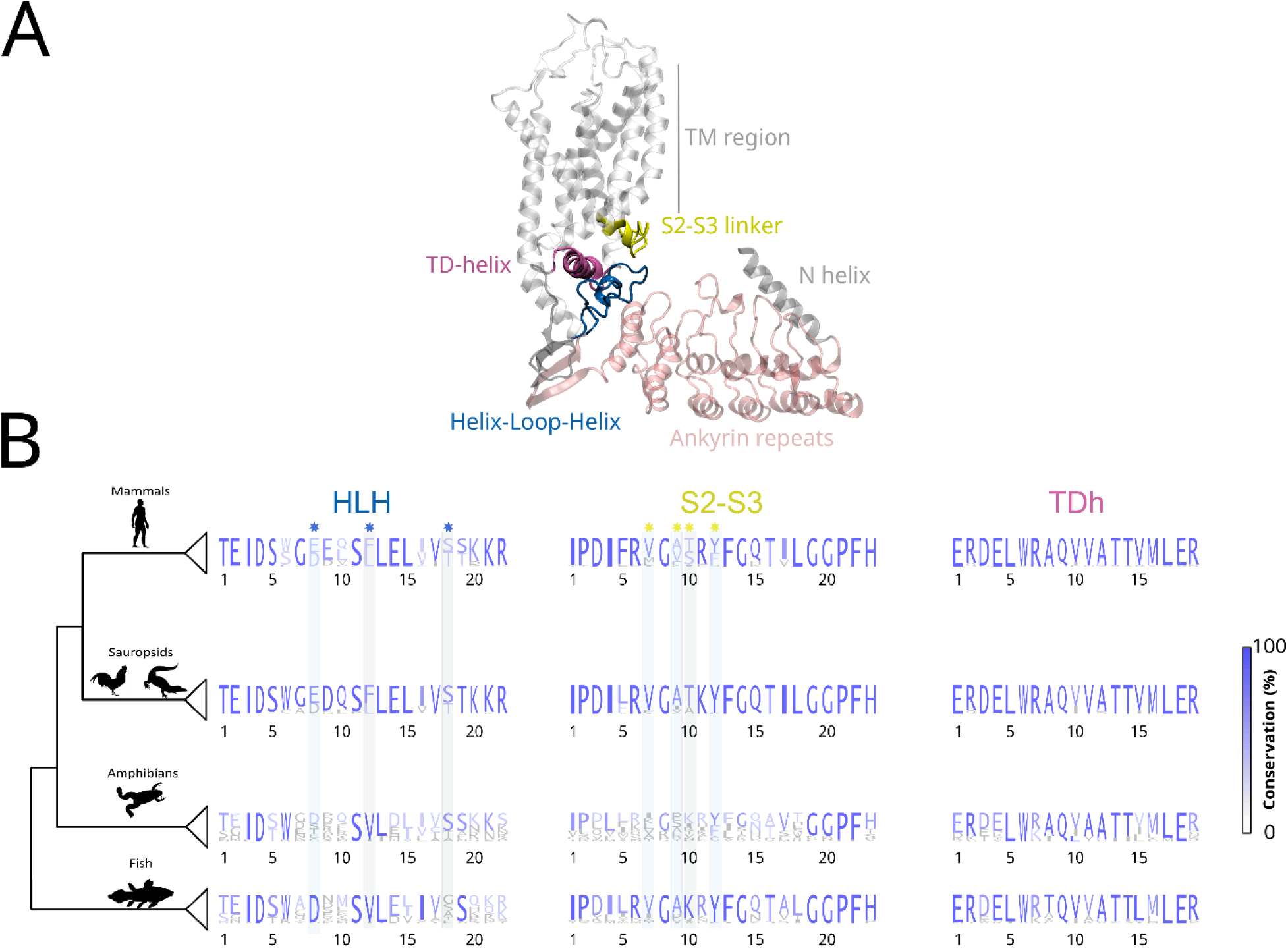
Helix-loop-helix, S2-S3 linker, and TRP domain helix domains show independent evolution patterns. A) Structure of one subunit of the human TRPV6 channel (hTRPV6, PDB:7S88). Ankyrin repeats domain (light pink), Helix-Loop-Helix domain (blue), S2-S3 linker (yellow), and TD helix (mauve) are highlighted. TM region: transmembrane region N helix: Amino terminal helix. B) Residue conservation (in size and blue shades) for the components of the HLH/S2-S3 linker/TDh inactivation motif. Shadows represent functionally relevant amino acids not conserved (gray) and conserved (blue) in amphibians and/or fish compared to sauropsids and mammals. Multiple sequence alignment in Supplementary figure 1.1 was used to calculate the percentage of amino acid identity. HLH: Helix-Loop-Helix domain, S2-S3 linker: linker between the transmembrane segments 2 and 3, TDh: TRP Domain helix.

In a previous study, we reported independent duplication events occurring in mammals and sauropsids (24). In both cases this duplication originates an evolutionary innovation in the form of a fast-inactivating channel ortholog. Here, our sequence analysis revealed that in mammals and sauropsids, the HLH domain has a scaffold of well conserved amino acids around the residues shown to be critical for fast inactivation (Fig. 1B, blue asterisks). Interestingly, such conserved scaffold is absent in amphibians and fish, whose Ca^2+^-selective TRP channels lack fast inactivation. Additionally, critical residues that are a fingerprint for fast-inactivating TRPV6 channels (i.e., Leucine at position 12 and Threonine at position 18, gray shadows) are not conserved in amphibian and fish, suggesting that these TRPV5-6 homologs have never evolved towards a fast-inactivating phenotype (Fig. 1B).

On the other hand, a similar but less strong pattern is detectable at the S2-S3 linker. Here, mammals and sauropsids have a more conserved set of amino acids around the residues defining the phenotype of fast inactivation (Fig.1B, yellow asterisks). Moreover, a threonine residue at position 10 (gray shadow) is highly conserved in fast inactivating TRPV6 channels and it is substituted by a well conserved positively charged residue in amphibian and fish TRPV5/6. However, three S2-S3 linker amino acids previously associated with inactivation kinetics in mammalian TRPV5 channels (Valine at position 7, alanine at position 9, and tyrosine at position 12, blue shadows) are conserved in fish. All this suggests that scaffolds of well conserved residues at the HLH and S2-S3 linker domains are required although not sufficient to support fast inactivation, and that the necessary fit between these two elements (HLH and S2-S3 linker) was not fully developed until the duplication event occurring at the mammalian ancestor.

Evolutionary reconstruction of the Transient Receptor Potential Vanilloid (TRPV) subfamily showed that calcium-selective TRPV5-6 and temperature-sensitive TRPV1-4 form independent monophyletic groups sister to each other (24). Our sequence analysis shows noticeable differences in the primary structure between these groups at the region of interest. Interestingly, inclusion of hTRPV1-4 channels, lacking Ca^2+^ dependent fast inactivation, introduced gaps in our alignments at both the HLH and S2-S3 linker (Supplementary Figure 1.1, zoomed regions black arrows). Sequence analysis showed significant divergence in the amino acid composition of the S2-S3 linker between the analyzed monophyletic groups. When comparing residues located at the HLH, we observed higher similarity between hTRPV1-4 and fish TRPV5/6 compared to the mammalian calcium-selective TRP homologs. Residues showed to modulate inactivation kinetics in mammalian TRPV5-6 (24) are not conserved in hTRPV1-4 (Supplementary Figure 1, black stars). This finding suggests that, similarly to amphibian and fish TRPV5/6, hTRPV1-4 did not evolve towards a fast Ca^2+^-dependent current decay mechanism.

Taken together, our sequence analyses suggest that molecular evolution associated to fast inactivation in TRPV5-6 mammalian channels was built on the stabilization of a conserved scaffold sequence surrounding specific fast inactivation-related residues that are located at the HLH and the S2-S3 linker.

### The distance between the HLH and TDh modulates fast inactivation

In the absence of a detailed inactivation mechanism, we worked under the underlying assumption that both phases of inactivation, fast and slow, converge into a structurally similar fully inactivated state. Accordingly, the apo/lipid-bound condition, defined by the absence of a bound ligand -other than lipids used for protein purification and imaging-was considered a non-inactivated state. Moreover, the Ca^2+^/CaM-bound structures were assumed as fully inactivated channels. To explore structural rearrangements occurred during inactivation, we compared the available structures of mammalian TRPV5 and TRPV6 in the apo/lipid-bound (PDB IDs for TRPV5: 6DMR, 6OP1; PDB IDs for TRPV6: 6BO8, 6BO9, 6BOB 7K4A, 7S88, 7S89) and Ca^2+^/CaM-bound states (PDB IDs for TRPV5: 6DMW, 6O20; PDB IDs for TRPV6: 6E2F, 6E2G).

To analyze the interactions within the inactivation motif, we measured the distance between residues located at the different domains (i.e., charged side chains putatively involved in interdomain salt bridges) in the available non-inactivated and fully inactivated TRPV5-6 structures. Average distances for each condition were used as a proxy for changes in interdomain interactions (Fig 2B). This close inspection revealed a specific set of residues that are in close proximity, suggesting they contribute to interdomain association (Fig 2A).

**Figure 2.**
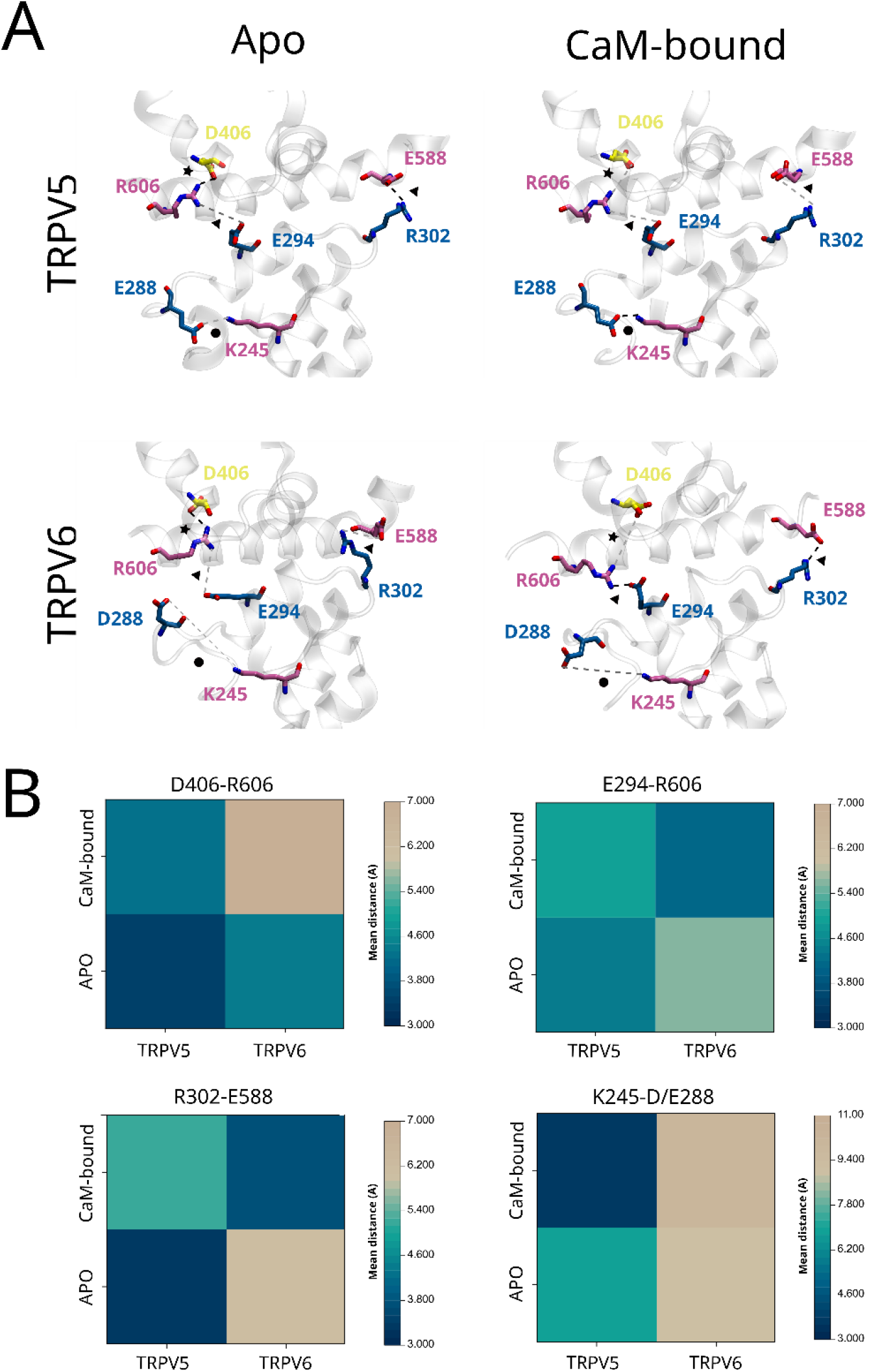
Evolutionary amino acid substitutions modulate the three-dimensional conformation of the HLH/S3-S3linker/TDh structural triad in mammalian TRPV5 and TRPV6. A) Structural arrangement of the three-dimensional triad of mammalian TRPV5 and TRPV6 in the apo/lipid-bound and Ca^2+^/CaM-bound conformations (PDBs; TRPV5 apo: 6DMR, TRPV5 Ca^2+^/CaM-bound: 6DMW, TRPV6 apo: 7S88, TRPV6 Ca^2+^/CaM bound: 6E2F). Residues located at HLH (blue), S2-S3 linker (yellow), TDh (mauve), and ARD (pink) contributing to interdomain interactions are highlighted. Interactions between HLH-TDh (triangle), HLH-ARD (circle) and TDh/S2-S3 linker (star) are highlighted. B) Heat maps representing the average distance between the pairs of residues in panel A. Bars on the right of each panel represent the color scale of the mean distance in the analyzed structures.

In non-inactivated TRPV5 channels, the S2-S3 linker is closer to the TDh when compared to TRPV6 (i.e., averaged distance between D406 and R606, star in Fig 2A) (Fig 2B). This observation extends to all other interdomain interactions including the HLH with respect to the TDh (E294-R606 and R302-E588, triangle in Fig 2A), and HLH to the ankyrin repeat domain 6 (ARD6) (E288-K245, circle in Fig 2A) (Fig 2B). In TRPV5, the majority of these interdomain distances increased when Ca^2+^/CaM complex is bound. Notably, the distance between HLH to ARD6 (E288-K245) is clearly shorter in the fully inactivated TRPV5 when compared to the apo structure, and to any conformation available for TRPV6 (Fig 2B).

The binding of the Ca^2+^/CaM complex to TRPV6 had an opposite effect on the interdomain distances. While residues located at the S2-S3 linker were farther from the TDh (i.e., D406-R606), we observed the TDh and the HLH domain remained in close contact (i.e., E294-R606 and R302-E588) (Fig 2B). In the fully inactivated TRPV6 channels the proximity between the HLH and TDh seems to be stabilized by the interaction between R302-E588 and E294-R606. We reasoned that the latter interaction prevents the closer contact between R606 and D406 (i.e., TDh and S2-S3 linker), observed in Ca^2+^/CaM-bound TRPV5, placing the HLH and TDh close to each other and away from the S2-S3 linker. In all conformations analyzed for TRPV6 the HLH and ARD6 are found apart from each other (Fig 2B).

The TRP domain helix runs parallel to the plasma membrane and is directly connected to the channel’s internal gate (Fig 3A). Thus, it is reasonable to think that the TDh drives the inactivation by coupling changes occurring in the landscape of interactions between the HLH and S2-S3 linker after binding Ca^2+^/CaM (Fig 3A). In this context, changes occurring to the association between the TDh to the HLH and S2-S3 linker would modulate the inactivation kinetics.

**Figure 3.**
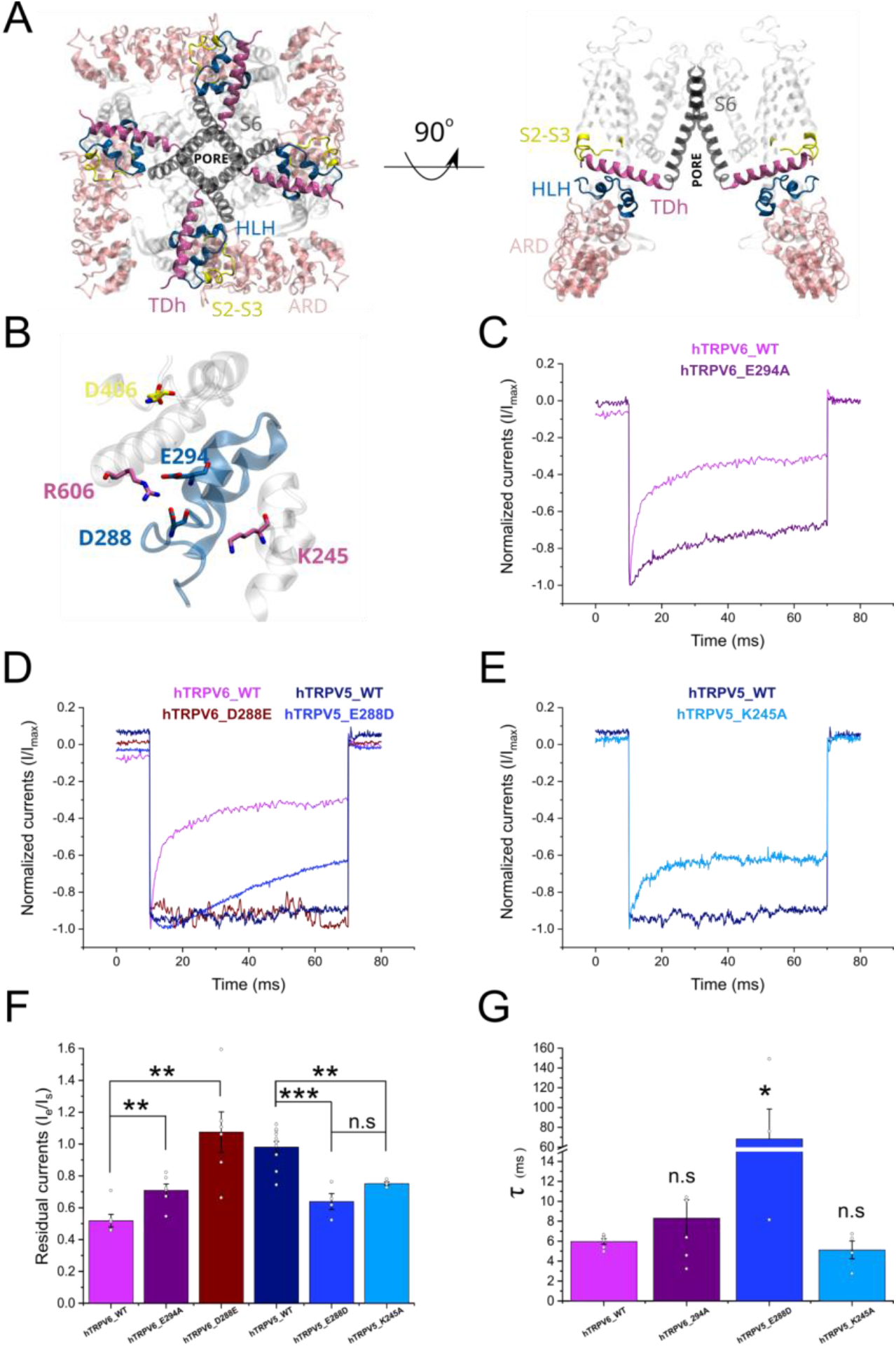
HLH interaction network modulates inactivation kinetics of hTRPV5 and hTRPV6 ion channels. A) Structure of the tetrameric hTRPV6 ion channel. The structure shows the TDh (mauve) connecting the S6 segment (dark gray) with the HLH (blue) and S2-S3 linker (yellow) domains. ARD in close contact with the HLH is shown in pink. Right panel shows two subunits for easier representation. B) Representation of residues involved in the differential interdomain interactions of the HLH in hTRPV5 and hTRPV6 (numbers according TRPV6). C) Normalized whole-cell current traces recorded from transiently transfected HEK-293T cells expressing wild type (WT) hTRPV6 and the mutant hTRPV6-E294A. D) Normalized whole-cell current traces recorded from transiently transfected HEK-293T cells expressing WT hTRPV5, hTRPV6, and the mutants hTRPV6-D288E and hTRPV5-E288D. E) Normalized whole-cell current traces recorded from transiently transfected HEK-293T cells expressing hTRPV5_WT and the mutant hTRPV5-K245A. F) Pooled data comparing the residual currents (defined as the ratio between the currents at the end versus the beginning of the pulse). G) Pooled data for the time constants of inactivation for clones showing a fast current decay. All current traces were recorded with 2 mM extracellular Ca^2+^ and elicited in response to a 60 ms pulse at −160 mV from Vh=0 mV (Panels C; D; E). Bars represent mean value; errors represent S.E. and white dots represent values for each experiment (Panels F; G). *** represents p = 0.001; **p = 0.01; *p = 0.05.

To test this, we first disrupted the interaction between the HLH and TDh in hTRPV6, by neutralizing the E294 negative charge. Patch-clamp recordings in whole-cell configuration showed that hTRPV6-E294A mutation partially prevents fast inactivation in hTRPV6 when compared to the wild type (WT) channel (Fig 3 C, F, G). The interacting E294 and R606 are highly conserved amino acids in TRPV5 and TRPV6, part of what we defined as scaffold sequences (Fig 1B positions 14 and 19 in HLH and TDh panels respectively).

Sequence and structural analysis highlight the amino acid at position 288 (Fig 1B position 8 in HLH panel), which is part of the previously described inactivation signature (24). While in TRPV6 this position is occupied by an aspartate exposed to the solvent, in TRPV5 it is a glutamate side chain that is in close contact with the K245 (located at the ARD6) in both the non-inactivated and fully inactivated states (Fig 2). Even though aspartate and glutamate are negatively charged amino acids, they differ in their side-chain length. This simple variation in length could be the molecular basis that modulates interdomain interactions required for inactivation. To test this idea, we exchanged D288 that is present in hTRPV6 with E288 of hTRPV5. Whole cell patch clamp recordings showed that the amino acid swapping caused a complete exchange of the fast inactivation phenotype (Fig 3 D, F, G). While hTRPV5-E288D showed a rapid current decay after activation, hTRPV6-D288E did not inactivate during the recording time. These results suggest that an interaction between the HLH and the ARD6 (i.e., E288-K245 in TRPV5) might stabilize the open non-inactivated state of the channel. To test this directly, we disrupted the putative electrostatic interaction by eliminating the K245 positive charge (hTRPV5-K245A). The hTRPV5-K245A mutant showed an evident fast inactivation that is absent in the WT counterpart (Fig 3 E-G).

Altogether, these results strongly suggest that close proximity between the HLH and the C-terminal region of the TDh (i.e., E294-R606) favors the fast inactivation observed in hTRPV6 channels. Moreover, this association is likely modulated by the interaction between critical residues located at the HLH with a positively charged residue located at the ARD6. We reasoned that this HLH-ARD interaction (i.e., E288-K245 in hTRPV5) might maintain the HLH away from the TDh favoring its contact with the S2-S3 linker (i.e., R606-D406 in hTRPV5) preventing fast inactivation in TRPV5 channels.

### Molecular dynamics simulations predict Ca^2+^ ions promote structural changes within the inactivation motif

So far, our analyses suggest that the conformational differences between the apo and fully inactivated TRPV5 and TRPV6 channels have a critical role in controlling the kinetics of the fast inactivation phase. Therefore, we reasoned that several of the structural rearrangements observed in the fully inactivated channels must occur during the fast phase of the inactivation process.

It has been proposed that the fast inactivation is triggered by Ca^2+^ ions in a concentration dependent fashion, suggesting direct binding (26). To explore whether the structural conformation of the HLH domain in the fully inactivated channels associates to Ca^2+^ ions, we performed short (100ns) fully atomistic molecular dynamics simulations (MDS) (Figure 4 and Supplementary Figure 4.1) in presence of 150mM NaCl or 50mM CaCl_2_ using apo-state structures (PDBs: 6DMR for TRPV5 and 6BOB for TRPV6). Interestingly, when analyzing the changes Ca^2+^ ions caused in the interaction network (Fig 4) we observed a very similar set of interactions described in the structural analysis (Fig 2). Ca^2+^ ions decreased the mean distance between the HLH and the C-terminal portion of the TDh (i.e., E294-R606, triangle) in TRPV6 compared to TRPV5 (Fig 4). While in TRPV5, Ca^2+^ binding brings the HLH and ARD (circle) closer, these residues move apart in TRPV6 (Fig 4). These results suggest that some of the interactions observed in the fully inactivated channel are likely catalyzed by calcium ions binding to the region during the rapid phase of the inactivation process.

**Figure 4.**
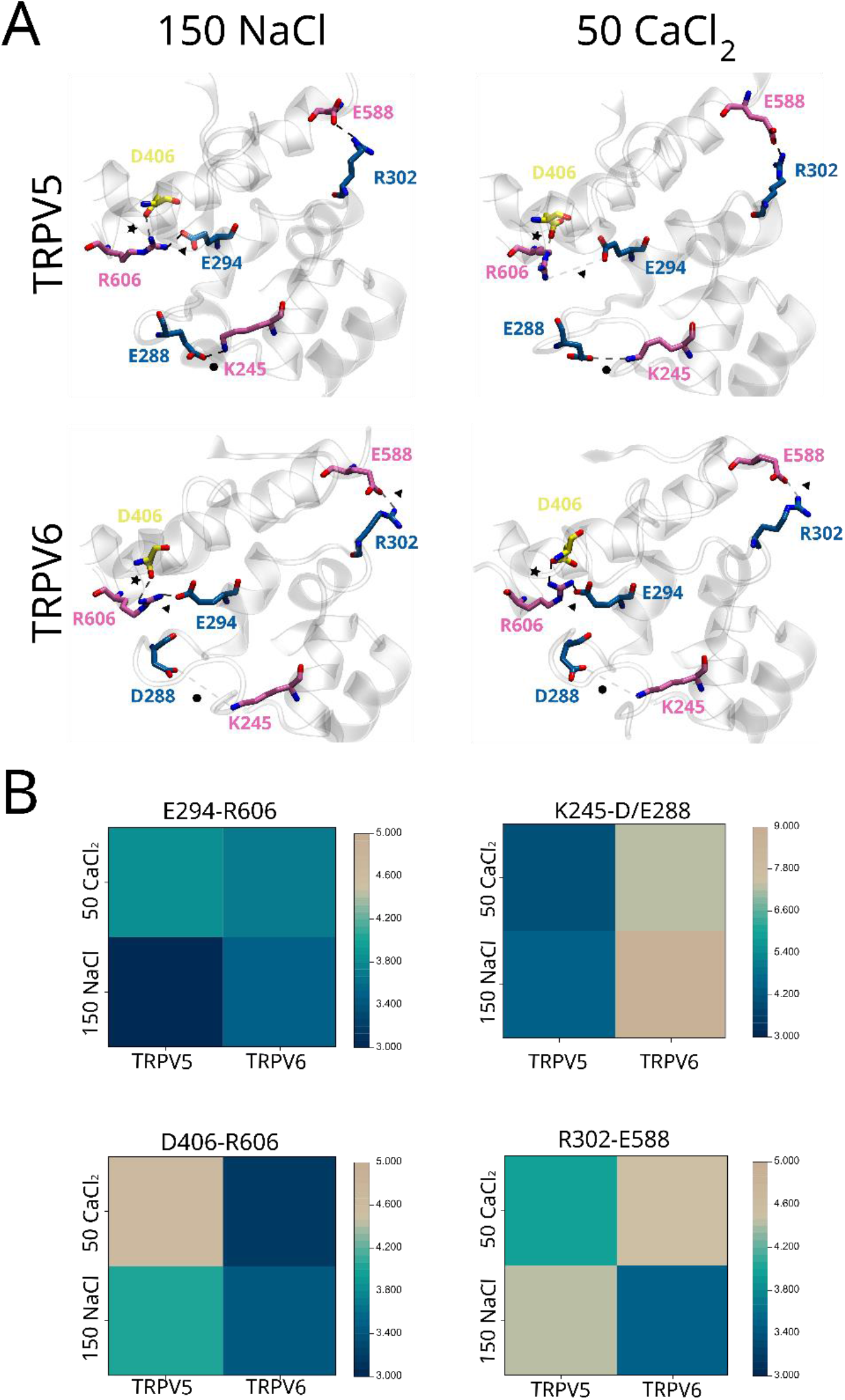
Calcium ions trigger structural changes within the HLH/S2-S3 linker/TDh inactivation motif in mammalian TRPV5 and TRPV6 channels. A) Structural conformation of the three-dimensional triad HLH/S2-S3 linker/TDh of mammalian TRPV5 and TRPV6 after running 100ns of full atomistic molecular dynamics simulations in presence of 150 mM NaCl or 50 mM CaCl_2_ (PDBs; TRPV5: 6DMR, TRPV6: 6BOB). Residues of the HLH (blue), S2-S3 linker (yellow), TDh (mauve), and ARD (pink) contributing to interdomain interactions are highlighted, as well as the interactions (HLH/ARD: circle, HLH/TDh: triangle, S2-S3 linker: star). B) Heat maps representing the average distance between the pairs of residues in panel A during the molecular dynamic simulations in presence of 150 mM of NaCl or 50 mM of CaCl_2_. Bars on the right of each panel represent the color scale of the mean distance during the simulation time.

We rationalized that the structural changes occurred during the fast and slow phases contribute to a unique inactivation mechanism. Therefore, we hypothesized that the structural arrangements common between the Ca^2+^ (i.e., result of molecular dynamics simulations) and the Ca^2+^/CaM-bound (i.e., fully inactivated structural models) states occur during the fast phase, while the structural differences observed are rather attributable to conformational changes occurring during the slow phase. In this context, the structural changes between the channels in presence of Na^+^ or Ca^2+^ are enhanced in the fully inactivated conformation. While in hTRPV6 the HLH and TDh (E294-R606) are even closer in the fully inactivated channel, their distance increased in hTRPV5. Enhanced differences were also observed when comparing the distance between HLH and ARD6 (i.e., E288-K245), with an increased proximity in hTRPV5. Our analysis suggests to us a set of consecutive conformational changes in the same direction, supporting the idea that the structural changes during the slow inactivation phase occur once the fast inactivation has taken place (Figure 5).

**Figure 5.**
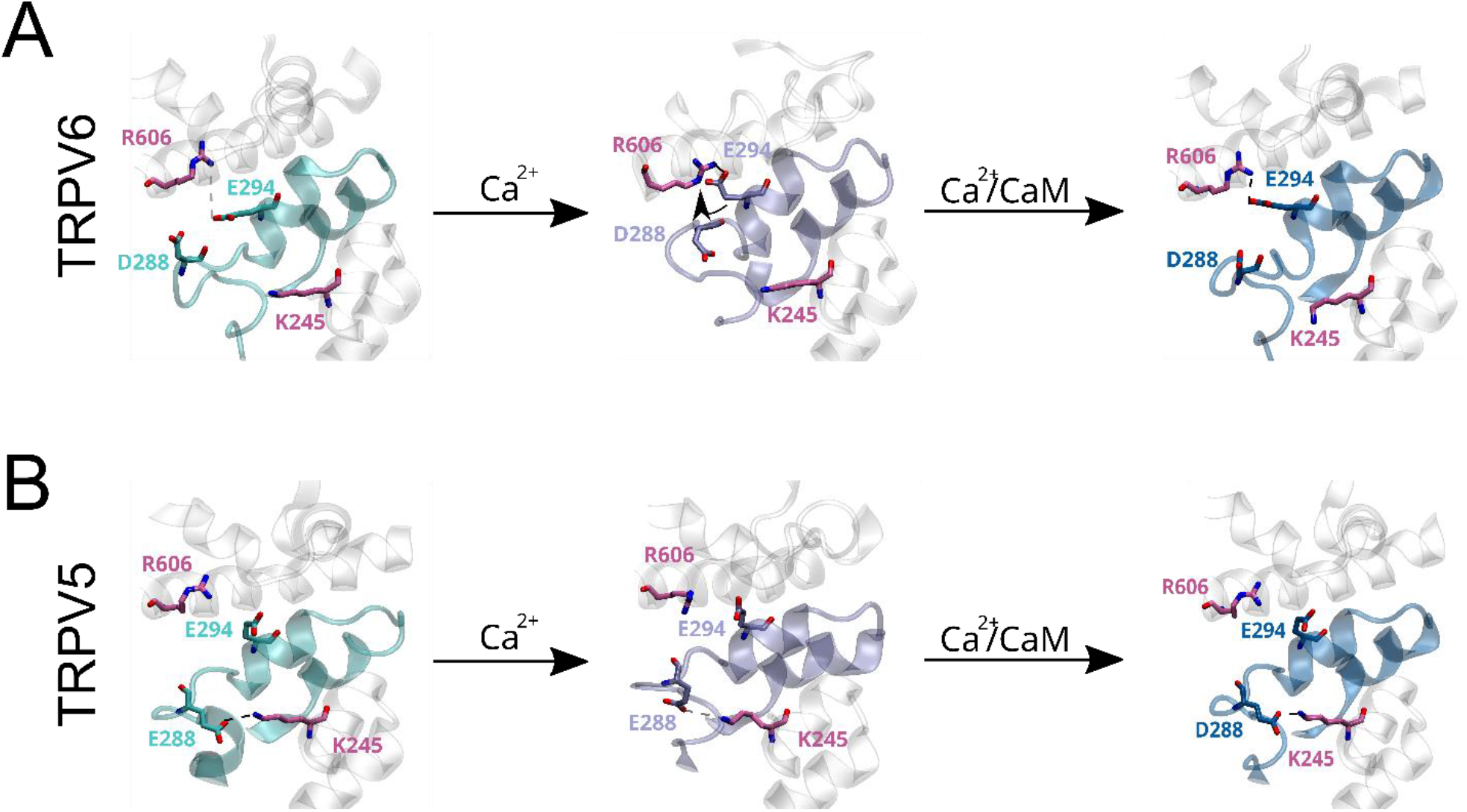
Proposed mechanism of fast inactivation modulation by the HLH in mammalian TRPV6 channels. Conformational changes induced by Ca^2+^ ions and the binding of the Ca^2+^/CaM complex to mammalian TRPV5 and TRPV6 channels. (A) Structural rearrangements bring closer the HLH (i.e., E294, blue) to the TDh (i.e., R606, mauve) in TRPV6. In mammalian TRPV5, an interaction between the HLH (E288, blue) and the ARD (K245, mauve), remains constant in every state, while no association between the HLH and TDh is observed.

## Discussion

### Possible mechanism of inactivation modulation by the HLH domain

Altogether, our evolutionary, structural, and functional analyses suggest that HLH is critical for the modulation of inactivation kinetics in mammalian TRPV5 and TRPV6 and that both inactivation mechanisms, fast and slow, likely share structural intermediates.

Our results suggest that molecular evolution tuned conformational changes occurring during the fast and slow inactivation phases. Based on our structural and functional analyses we propose a sequential model for the changes occurring during Ca^2+^-dependent inactivation. Briefly, once the channel opens, Ca^2+^ ions permeate through the pore increasing its intracellular concentration. Elevated local [Ca^2+^]_i_ at the channel intracellular microenvironment disrupts the association between the TDh and the S2-S3 linker bringing the HLH and TDh closer. Finally, the structural rearrangements occurring at the C-terminal region of the TDh are transmitted to the pore though the same helix whose N-terminal region is physically connected to the channel internal gate (Figure 5A). In this context, our results suggest that the interaction of the residue 288 at the HLH with the ARD6 (i.e., K245) is critical in defining the positioning of the HLH by holding it back, preventing its interaction with the TDh and hence fast inactivation. Proximity between HLH and the TDh is easier to achieve in TRPV6 just because the interaction between HLH and ARD6 is absent. We reasoned that the strong association between the amino acids E288 and K245 in all the conformational states of mammalian TRPV5 increases the energetic barrier for the HLH to interact with the TDh, and therefore prevents the rapid current decay (Fig 5B). The closer proximity between HLH and TDh in TRPV6, and HLH and ARD in TRPV5 are even more evident in the fully inactivated channels suggesting a common ground to both inactivation mechanisms.

### Molecular evolution defines biophysical properties of TRPV channels

Our multiple sequence alignment of members of the TRPV subfamily showed a high level of conservation (Supplementary Figure 1.1), especially in the TDh, as expected for *bona fide* TRP channels. Nevertheless, some sequence gaps are observed between the sister monophyletic groups TRPV1-4 (thermoTRPV) and TRPV5-6 (calcium-selective TRP). Interestingly these gaps are in highly relevant functional domains. Gaps 1 and 2 are located at the ARD and HLH domains. Residues located in these regions have been related to the temperature-dependent activation of TRPV1 and TRPV3 (27–30), while our work suggest the two domains have a significant role in modulating Ca^2+^-dependent fast inactivation, a biophysical property unique to TRPV5-6. The third gap is in the S2-S3 linker. To the best of our knowledge this domain is not involved in modulating the activation by temperature of TRPV1-4, but previous works have shown this is a key modulator of the fast inactivation in TRPV5-6 (23,24). Also, in other TRP channels the S2-S3 linker binds Ca^2+^, but residues important for such coordination are absent in the TRPV subfamily (31) and the cation binding was not observed in our molecular dynamic simulations; studies addressing the Ca^2+^ binding process in TRPV5-6 are required.

Here, we propose that the scaffold of highly conserved amino acids located at the HLH and S2-S3 linker of mammalian TRP5-6 channels, supports the three-dimensional structure required for the correct complementarity between the elements forming the inactivation motif. These scaffold sequences, absent in amphibians and fish TRPV5/6 and hTRPV1-4, were probably generated at the common ancestor of mammalian and sauropsids TRPV5-6, while specific amino acid modifications occurred only in one of the duplicated genes in mammals, resulting in fast inactivation as an evolutionary innovation.

Multiple groups have reported the relationship between the molecular evolution and functional properties of the superfamily of TRP channels, but not much is known about the evolution of TRPV5-6 functional properties. Altogether our results suggest that the molecular evolution within the TRPV subfamily led to the unique functional properties of the sister TRPV1-4 and TRPV5-6 monophyletic groups. While TRPV1-4 evolved towards a temperature-dependent mechanism of gating, molecular evolution of TRPV5-6 resulted in inactivating channels that further evolved a simple mechanism for fast inactivation as an evolutionary functional innovation.

## Methods

### Sequence data and analysis

We retrieved calcium-selective TRP channels sequences in representative species of all major groups of vertebrates. Our sampling included species from mammals, birds, reptiles, amphibians, coelacanths, holostean fish, teleost fish, cartilaginous fish, and cyclostomes (Supplementary Figure 1). Protein sequences were obtained from the Orthologous MAtrix project (OMA)(32). In cases where the species were not included in the OMA project, we searched in the NCBI database (refseq_genomes, htgs, and wgs) using tbalstn (33) with default settings. Protein sequences were aligned using the FFT-NS-1strategy from MAFFT v.745. (34). Human TRPV1, TRPV2, TRPV3, and TRPV4 were used as out groups.

### Molecular Biology, Cell Culture and Transfection

Open Reading Frames (ORF) codifying to the different channels analyzed [human TRPV5 (hTRPV5) WT, hTRPV5_E288D, hTRPV5_K245A, human TRPV6 (hTRPV6) WT, and hTRPV6_D288E], inserted in a pcDNA3.1(+) vector, were obtained from GenScript Corporation (Nanjing, China). HEK 293T cells were grown in DMEM-F12 medium containing 10% (v/v) bovine fetal serum at 37ºC in a humidity-controlled incubator with 5% (v/v) CO2. HEK 293T cells were transiently co-transfected with the different clones analyzed and peGFP-N1 to allow their identification.

### Electrophysiology and solutions

Whole-cell currents were measured with an Axopatch-200B amplifier. We used borosilicate pipettes (o.d. 1.5mm, i.d. 0.86mm, Warner Instruments, Hamden, CT) with resistance of 2 to 4.0MΩ to form seals with access resistance between 2 and 6 GΩ. Macroscopic currents were recorded in response to a voltage step protocol from zero to -160mV. Inactivation was analyzed in a time window of 60 ms. Recordings were digitized at10KHz and filtered at 5KHz using a Digitada 1320 (Molecular Devices, LLC, California). The analysis was performed using Clampfit 10.3 (Molecular Devices, LLC, California). Fast inactivation was assessed by computing the residual currents, defined as the ratio of the current value at the end of the negative pulse over the current at the beginning of the voltage pulse (ref). The standard extracellular solution (Ringer-Na^+^) contained (in mM) 140 NaCl, 5 KCl, 2 CaCl_2_, 2 MgCl_2_, 8 glucose and 10 HEPES, at pH 7.4 adjusted with NaOH. The standard internal (pipette) solution contained (in mM) 105 CsF, 35 NaCl, 10 EGTA and 10 HEPES, at ph 7.4 adjusted with CsOH. All the experiments were performed at room temperature (20 to 25ºC).

### Statistical Analysis

Data are expressed as the mean ± S.E. Overall statistical significance was determined by analysis of variance (ANOVA one way) with a Bonferroni post-test, and T-student tests. For all conditions, the average was obtained from at least 4 independent experiments. The outliers were defined by using Graphpad QuickCalcs (https://www.graphpad.com/quickcalcs/Grubbs1.cfm) and removed from the analysis.

### Structural analysis of the available structures of mammalian TRPV5-6 channels

Available three-dimensional structures of mammalian TRPV5 (PDB IDs: 6DMR, 6OP1, 6DMW, 6O20) and TRPV6 (6BO8, 6BO9, 6BOB 7K4A, 7S88, 7S89, 6E2F, 6E2G) were used to study the conformational states of the HLH/S2-S3 linker/TDh in the apo/lipid-and Ca^2+^/CaM-bound states. To determine the distance between residues putatively involved in interdomain interactions were measured the distance between the last negatively charged oxygen and positively charged nitrogen in VMD 1.9.3. Resultant distance values for each condition were averaged and used as a proxy to study conformational differences.

### Molecular dynamic simulations (MDS)

Three-dimensional structures of rat TRPV6 (PDB: 6BOB) and rabbit TRPV5 (PDB: 6DMR) were prepared at pH 7.0 and subjected to energy minimization in vacuum through the Maestro-Schrödinger suite (Schrödinger Release 2018-2: Maestro, Schrödinger, LLC, New York, NY,2018). The channels were embedded into a pre-equilibrated POPC lipid bilayer in an orthorhombic box with periodic borders, filled with Single Point Charge (SPC) water molecules and an ionic concentration of 150 mM NaCl or 50mM CaCl_2_. Full atom Molecular dynamics simulations of 100ns were performed, per triplicates, in a NPT ensemble (P = 1 atm, T =310 K) with the Desmond software and OPLS v.2005 force field (35). Structures were collected every 0.2 nanoseconds during the MDS having 502 frames per simulation. Mean interaction distance between the residues was determined by averaging the measured distance between the last negatively charged oxygen and positively charged nitrogen along the simulation time in VMD 1.9.3.

## Acknowledgements

This work was supported by Fondo Nacional de Desarrollo Científico y Tecnológico from Chile (FONDECYT 1191868) to SB, ANID-Millennium Science Initiative Program #NCN19_168 (SB), and Comisión Nacional de Investigación Cientifíca y Tecnológica (CONICYT 21150220) to LFA. The Millennium Nucleus of Ion Channel-Associated Diseases (MiNICAD) is a Millennium Nucleus of the Iniciativa Milenio, National Agency of Research and Development (ANID, Chile)

---

**Figure 1- Supplement Figure 1.**
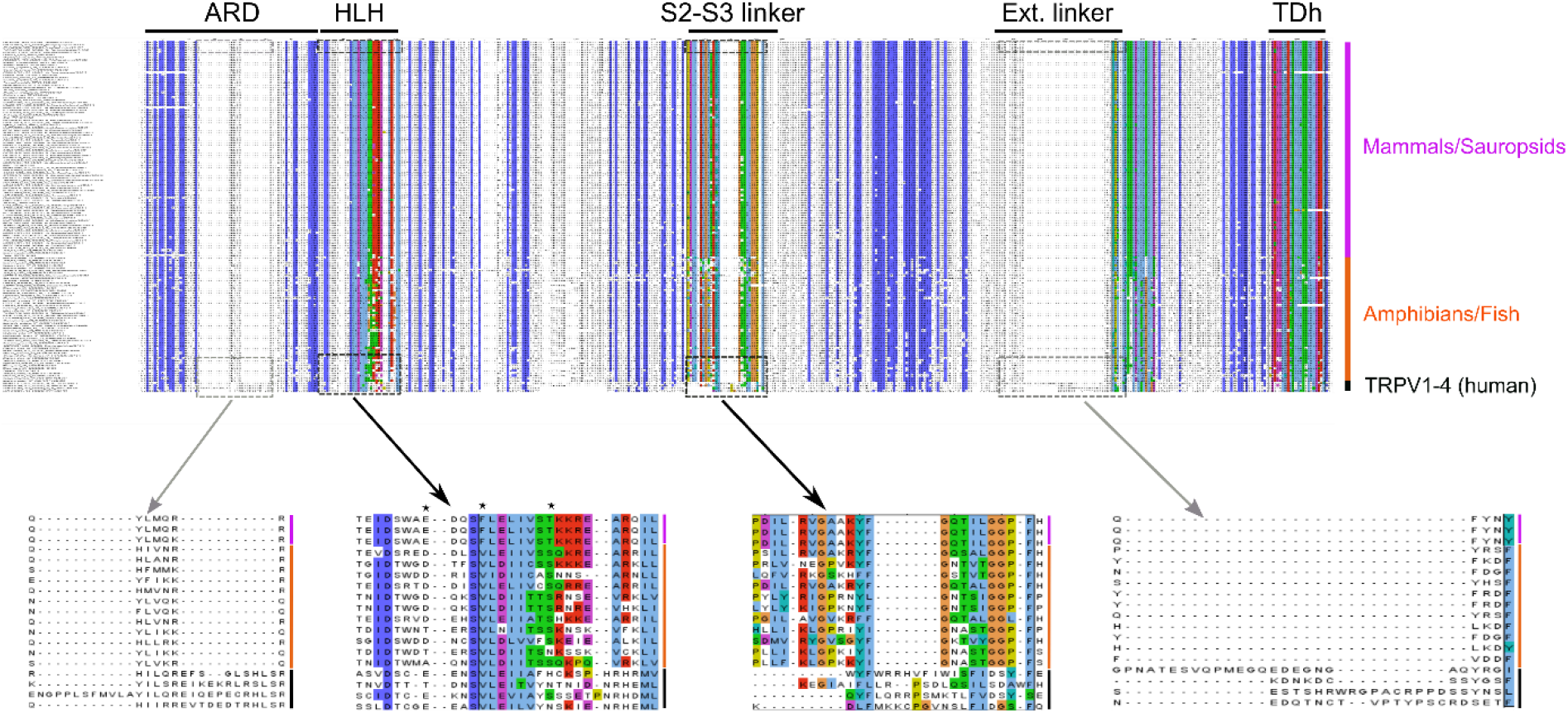
Multiple sequence alignment of TRPV5 and TRPV6 in vertebrate representative species. Sequences of hTRPV1-4 were used as an external group. Gene and species are stated to the left and the amino acid position within the alignment is in the central superior region. Different regions of the proteins are highlighted on the top as: ARD (ankyrin repeat domain), HLH (helix-loop-helix domain), S2-S3 linker (linker between the intracellular transmembrane segments 2 and 3), Ext. linker (extracellular linker known as turret in hTRPV1), TDh (TRP Domain helix). Sequences in zoom highlight the sequence gaps in the HLH, S2-S3 linker (black), ARD and extracellular linker (gray) between the two monophyletic groups. Color bars next to the zoomed alignment represent the group (mauve: mammalian, orange: fish, black: hTRPV1-4).

**Figure 4 – Supplement Figure 1.**
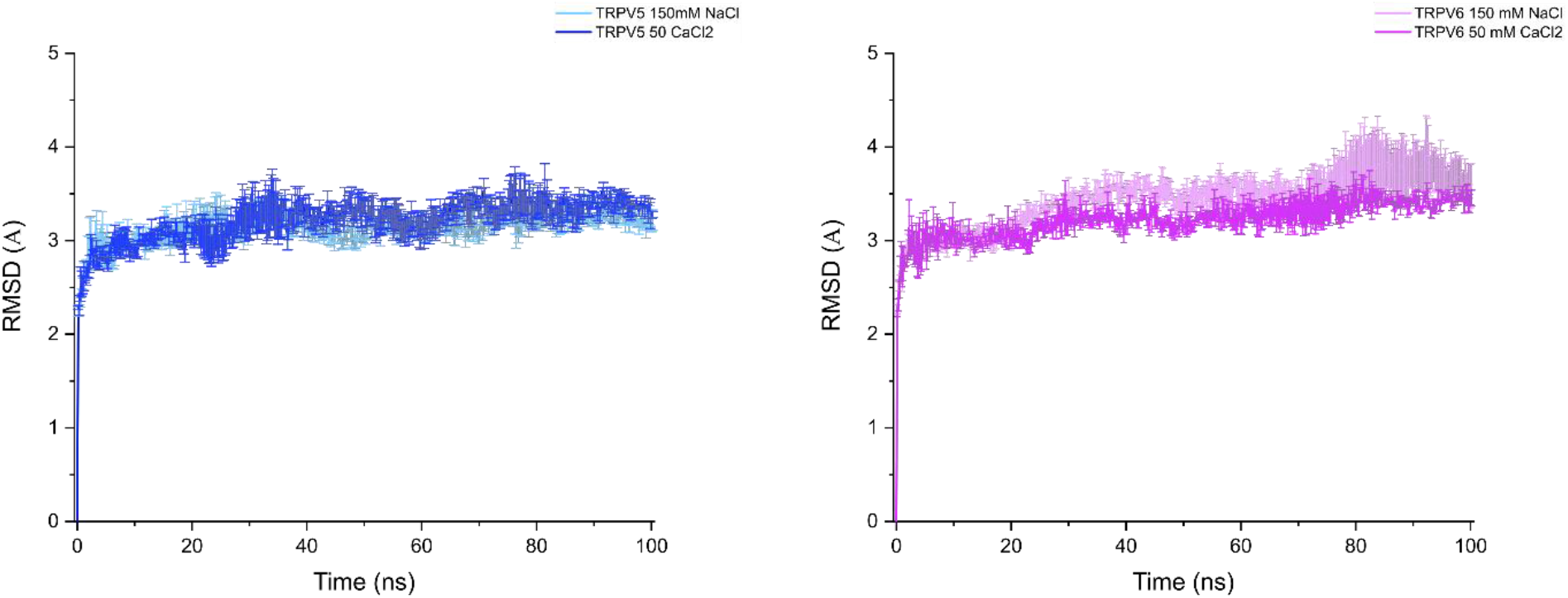
RMSD of TRPV5 and TRPV6 proteins during the 100 ns of molecular dynamic simulations in presence of 150 mM NaCl (light) or 50 mM CaCl_2_(dark).

